# 30S-seq redefines the bacterial Ribosome Binding Site

**DOI:** 10.1101/2025.04.04.647229

**Authors:** Jonathan Jagodnik, Valentin Gherdol-Nouvion, Mira Al Sahmarani, Alexandre Maes, Maude Guillier

## Abstract

The translation initiation step is rate limiting for the efficiency of gene expression in all organisms. However, the mechanism of ribosome recruitment to mRNA start sites strikingly differs between eukaryotes and prokaryotes. The eukaryotic small (40S) ribosomal subunit binds 5’ end caps and scans for the start codon while the bacterial small (30S) subunit directly binds to the Shine-Dalgarno (SD) motif close to the initiation site. Pioneer studies have shown rare 30S loading events further upstream within 5’ untranslated regions (5’UTRs), at ribosome standby sites^1–3^. Together with the frequent occurrence of long bacterial mRNA 5’UTRs and degenerated SD sequences, this indicates that the 30S subunit might bind upstream of the SD more commonly than currently thought. We therefore developed 30S-seq to map 30S-mRNA interactions in a bacterial transcriptome (*Escherichia coli*), inspired by translation complex profile sequencing (TCP-seq) previously used in eukaryotes^4,5^. Our results provide new and unsuspected insights into the behaviour of 30S and 70S complexes during the canonical translation initiation process. Notably, 30S subunits are recruited upstream of the start codon, primed to receive the SD released by the departing 70S ribosome. Remarkably, we also find hundreds of non-canonical 30S binding sites within mRNA 5’UTRs, sometimes over 100 nucleotides upstream of the start region. We validated several of these upstream ribosome binding sites, and demonstrated their strong impact on gene expression. Thus, even in bacteria, ribosomes frequently bind mRNAs outside of the start region to initiate translation, challenging the classic ribosome binding site model.

---

Translation initiation is a conserved process resulting in the assembly of a translation initiation complex around the mRNA start codon. Yet, the mechanisms by which the start codon is found strongly differ in eukarya and bacteria. The eukaryotic small subunit of the ribosome (40S) most often binds to the 5’ end of mRNAs through the 5’methyl-guanosine cap^6^. Upon docking to 5’ caps, the 40S subunit scans mRNAs for functional translation initiation codons. In striking contrast, the bacterial small subunit of the ribosome (30S) is mostly thought to directly bind to mRNA translation initiation regions (TIRs). This dogma originated from the discovery of the conserved sequence at the 3’ end of the 16S rRNA^7^. This region base pairs with its mRNA Shine-Dalgarno (SD) motif counterpart in mRNA^8^, typically located 4-12 nucleotides (nts) upstream of start codons. 30S binding to circularized mRNA templates later excluded any requirement for a free 5’ end^6^, further supporting a model where the 30S subunit binds directly to TIRs. The key role of the SD for gene expression was also confirmed in many studies ^9–11^.

Certain lines of evidence have left open the possibility of other modes of 30S subunit recruitment. Indeed, mRNAs carrying a poor SD or even lacking one entirely can be highly translated. This is the case for the SD-less *rpsA* mRNA encoding the ribosomal protein S1, one of the most abundant proteins in *Escherichia coli* (*E. coli)*^12^. In addition, mRNA features other than the SD or the start codon can promote 30S binding, such as secondary structures positioned immediately downstream of the TIR^13^. Such a structure is for instance present in the *bamA* mRNA, encoding an essential protein involved in the assembly of the outer membrane β-barrel proteins. Interestingly, these activating stem-loop structures are reminiscent of similar downstream structures facilitating the recognition of the start codon in eukaryotes^14^. Together with the absence of a recognizable SD motif in the *bamA* mRNA, this suggests that the 30S subunit could primarily bind to certain mRNAs in the 5’UTR and later translocate to the TIR. There, it would be stabilized by the translation activating stem-loop. To date, 30S binding to sites upstream of the TIR has been demonstrated in only a few rare examples^1–3^, including one in *E. coli* within the *tisB* toxin-coding mRNA^2,15^. There, the 30S subunit binds to a standby site, until the appropriate conditions (*e*.*g*., structural refolding of the TIR) lead to 30S translocation to the TIR for translation initiation.

Together with the poor conservation of the SD sequence in about 20% of mRNAs in Gammaproteobacteria^16^, these data highlight the possibility that the 30S subunit can bind outside of TIRs, but the extent to which such events occur has never been established. In eukarya, translation complex profile sequencing (TCPseq) revealed the different steps of translation initiation, including the binding of the 40S subunit to mRNAs^4,5^. This notably validated the predominant 5’cap-dependent scanning mechanism and the steps of full ribosome assembly at start codons. Instead, ribosome profiling approaches used in bacteria to study translation initiation have focused on the analysis of retapamulin-stalled 70S particles. While this method successfully established maps of translation start sites^17,18^, it did not provide details on 30S positioning at TIRs or the possible binding of the 30S subunit outside of TIRs. To address these questions, we developed 30S-seq and produced the first global map of 30S binding sites in the bacterial model *E. coli*. These data highlight both known and previously unknown conformations of the mRNA in complex with the 30S and 70S ribosomes, and provide an overview of the dynamics of ribosome binding and translation initiation. They furthermore reveal hundreds of 30S binding sites both distant from and close to the TIRs. We confirmed 30S binding to selected 5’UTRs, and showed that these sites have an extensive impact on gene expression. This study revisits our understanding of translation initiation in bacteria, even in a model as well-studied as *E. coli*.

## 30S-seq successfully identifies 30S- and 70S-binding sites in *E. coli*

The strategy of the 30S-seq technique is based on the TCP-seq approach that was developed in yeast^4,5^, with major experimental adjustments made for bacteria (Fig. 1a and Extended Data Fig. 1a). Briefly, *E. coli* cultures grown to exponential phase were snap-chilled in a Mg^2+^-rich ice to preserve ribosome-mRNA interactions. Cells were either immediately formaldehyde-crosslinked to further stabilize ribosomes at any stage of translation, or non-crosslinked as a reference. Translated RNAs carrying at least one 70S particle were sedimented and digested with micrococcal nuclease (MNase) to isolate 70S- and 30S-protected mRNAs regions (translated fraction-associated 30S, T-30S). The 30S fraction that sedimented separately from translated RNAs (Free-30S) was also isolated and MNase-treated. This Free-30S fraction contains 30S particles associated with mRNAs before the first round of translation starts or after the last round ends, in addition to unbound 30S. 15-45 nt-long ribosome-protected fragments were analyzed by Illumina high-throughput paired-end sequencing. Total RNAs and translated RNAs were also sequenced to provide input controls.

**Figure 1.**
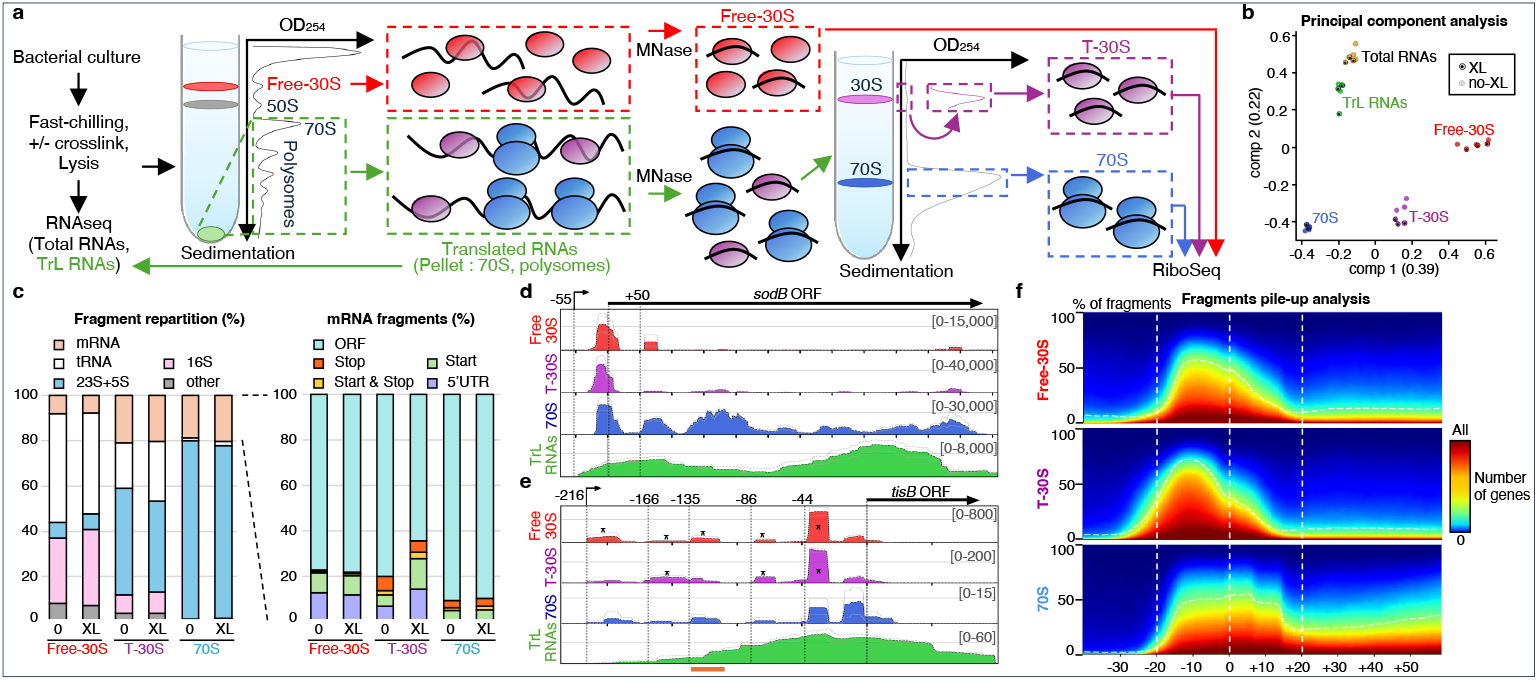
30S-seq reveals the details of 30S binding to TIRs and 5’UTRs. **a**. Outline of the 30S-seq. See text for details. **b**. Principal Component Analysis of all the 30S-seq and RNAseq samples (based on whole transcript FPKM levels). **c**. Repartition of protected fragments aligning to whole transcriptome (left panel) and various mRNA regions (right panel) in Free-30S, T-30S or 70S fractions in crosslinked (XL) or non-crosslinked (0) samples. Start and Stop correspond to fragments that contain the full codon, and Start-Stop fragments contain both a Stop codon and the Start codon of the subsequent gene, usually in a polycistronic mRNA. 5’UTR fragments align to mRNAs upstream of the third nt of the start codon. (**d, e**) 30S-seq profiles of *sodB* (**d**) and *tisB* (**e**) mRNAs. Positions relative to the translation start are displayed, and the open reading frame (ORF) and the transcription start site (TSS) are represented by arrows. The average of three biological replicates is shown, with “maximum” and “minimum” replicates in dotted lines. The pileup for each sample is normalized by the overall scaling factor. *: site detected as an uRBS in this study (see description below). The position of the previously demonstrated 30S binding site in *tisB* 5’UTR^2,15^ is underlined in orange. **f**. Global footprint of 30S-seq data on the 1919 first genes of operons with at least 15 fragments at the TIR (in at least one fraction). Positions relative to translation start are indicated. All genes are normalised to weight similarly in the analysis. The y axis represents the percentage of fragments relative to the total in the same area.

Multiple indicators clearly validate the 30S-seq approach and highlight strong differences between the Free-30S, T-30S and 70S fractions. These include principal component analysis (PCA), fragment sizes and distributions among the different fractions, as well as mRNA 30S-seq profiles (Fig. 1b-e, Extended Data Fig. 1b-g, 2 and Supplementary Information). In particular, the distribution of the reads among rRNAs, tRNAs and different regions of the mRNAs demonstrates that the three isolated fractions constitute distinct ribosomal populations, as expected. Of important note, the crosslink step had only a relatively mild impact on the overall count and distribution of the reads, reinforcing the biological relevance of the 30S-seq results. The crosslink step affected mostly the T-30S samples, where it led to a ∼2-fold enrichment in TIR- and 5’UTR-associated reads (Fig. 1c). This is likely due to a significant reduction of 70S dissociation during steps following crosslinking in the 30S-seq experiment (e.g., *sodB* and *dps*, Extended Data Fig. 1 c, d). Hence, the crosslinking step stabilized 70S particles and helped to isolate the T-30S ribosomal population of interest. For simplicity, we focused on crosslinked samples for the rest of the study.

As expected, multiple mRNAs displayed 30S-seq profiles in which the 30S subunits accumulated specifically in TIRs, while 70S ribosomes were found at TIRs and spread along the downstream ORF (*e*.*g*., *sodB* and *dps* mRNAs, Fig. 1d and in Extended Data Fig. 1c, d). The same was true when we assessed the global ribosome coverage at and adjacent to the TIRs of the first ORFs of operons, through a metagene analysis (Fig. 1f), further validating the technique. We also retrieved known internal translation start sites, such as the one recently described within the *arcB* ORF^17^ (Extended Data Fig. 1e). Finally, we found clear 30S binding sites within the *tisB* 5’UTR, that are most enriched in the Free-30S fraction (Fig. 1e). This includes the well-described standby site at position -135 relative to the translation start^2,15^. Importantly, 70S particles were only present at background levels at these sites, excluding the possibility that 30S binding took place at unknown translation initiation sites or resulted from the dissociation of elongating 70S ribosomes.

Hence, the 30S-seq successfully mapped 30S subunits and 70S ribosomes at expected sites within mRNAs, and appears perfectly suited for the transcriptome-wide study of ribosome binding and translation initiation.

### 30S-seq reveals new translation initiation intermediates

We noticed that the Free-30S and T-30S binding patterns at TIRs were different, *e*.*g*., in the individual 30S-seq profiles such as for *sodB, dps* or the internal start site within *arcB*, with T-30S accumulation shifted upstream of the start codon (Fig. 1d and Extended Data Fig. 1c-e). Importantly, the same trend was visible with the global ribosome coverage analysis (Fig. 1f). Indeed, T-30S subunits accumulated predominantly upstream of the translation start, while the Free-30S and 70S were more centered around the start codon. To obtain greater detail about the precise 30S and 70S positioning and sequence of events during translation initiation, we created a map of ribosome occupancy by analyzing the 5’- and 3’ end positions of each fragment in the (−40+40) region (Fig. 2).

**Figure 2.**
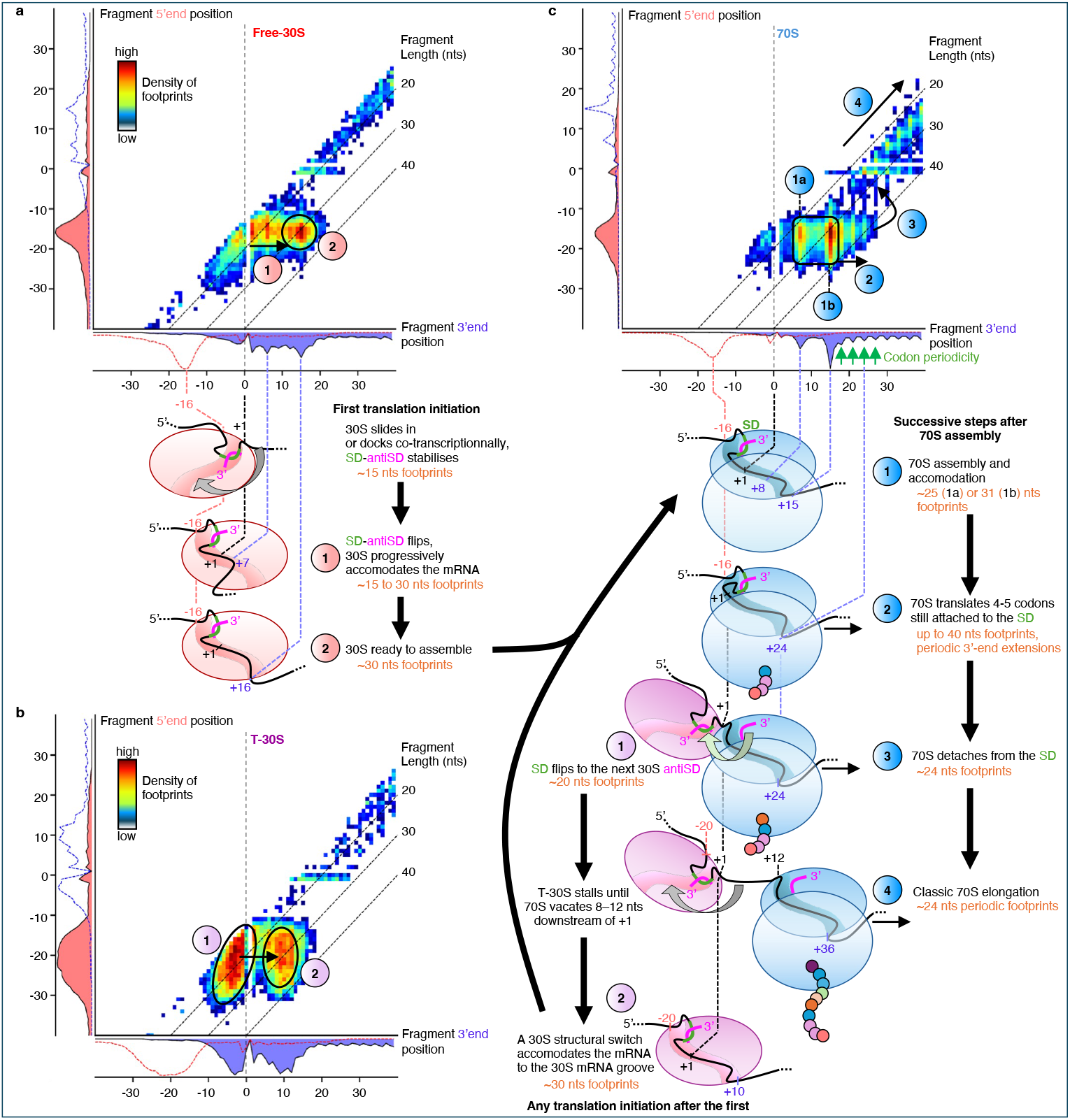
30S-seq recapitulates the dynamics of 30S accommodation and translation initiation in *E. coli*. Heatmap of Free-30S (**a)**, T-30S (**b**) and 70S (**c**) extremities of protected fragments in a subset of 1919 genes that are first in their operons, weighted by the overall per-gene extremity’s density. For each footprint, the position relative to the translation start of the 5’ end is plotted against that of the 3’ end. One-dimensional heatmap projections of 5’ ends and 3’ ends are represented in red and blue, with their 3’ and 5’ projection counterparts in dotted lines, respectively. The codon periodicity is indicated by green arrows on the x-axis for 70S fractions. Hotspots accumulating the most fragments (hottest pixel colour) are labelled and interpreted in the model, with the corresponding footprint sizes in orange font. Further description of the model is provided in the text. Note the expected bias caused by MNase digestion, at position +2 (usually 5’ of a G), causing a stark diminution of fragments ending at this position. The same representation for the non-crosslinked samples is shown in Extended Data Fig. 3.

Again, the patterns were different in the Free-30S, T-30S and 70S fractions. As expected, the signals for the Free-30S and T-30S were mostly around the TIR, with major footprints well-positioned for translation initiation, covering the region (−15+15) for the Free-30S (conformation 2 in Fig. 2a) and (−20+10) for the T-30S sample (conformation 2 in Fig. 2b).

In addition, some footprints present in both 30S fractions showed the existence of other conformations that differed in the translated or non-translated fractions. Indeed, the Free-30S fraction displayed rather continuous footprints with a 5’ end around -15 and a 3’ end extending roughly from -1 to +15. These likely correspond to the transition from early 30S-mRNA complexes to an accommodated state, ready for 70S assembly (conformation 1 in Fig. 2a).

In contrast, a second major footprint in the T-30S fraction was centered around the (−20-5) region. Hence, most protected fragments in this region did not include the start codon (conformation 1 in Fig. 2b). This bimodal distribution suggests a rapid switch from conformation 1 to 2 that contrasts with the continuous accommodation in the Free-30S sample. As this is only observed with the T-30S sample, it is likely related to the translation process. We thus analyzed the occupancies within the 70S fraction (Fig. 2c). This revealed several groups of footprints covering the SD and start codon (5’ end centered on -15 and 3’ end extending from +8 to +25). Strikingly, the codon periodicity was already visible from the 3’ end starting at position +15. This suggests that translation of the first codons takes place with the 70S still anchored at the SD (conformation 2 in Fig. 2c). We then observed fragments of a constant length of 24 nts following a codon periodicity, only with 5’ ends at or downstream of +1. These fragments are typical of translation elongation, and suggest that 70S ribosomes at this position are no longer interacting with the SD (conformation 3 in Fig. 2c). Together, the patterns of the T-30S and 70S conformations suggest that upon release of the interaction with the 70S ribosome, the SD flips to the next 30S subunit, primed to initiate translation behind the departing 70S ribosome (conformation 1 in Fig. 2b). Once the 70S ribosome liberates the early coding region, the 30S subunit accommodates this region to the mRNA tunnel (conformation 2 in Fig. 2b). It is then in the correct conformation for 70S assembly and the next translation cycle.

### Strong SD motifs correlate with 30S subunits primed upstream of the start codon

We wished to further understand the contribution of the T-30S occupancy upstream of the start codon to the translation initiation process. We therefore quantified the enrichment (ribosome coverage normalized to RNA levels) for the first genes of operons in various mRNA regions: upstream of nt -20 in the 5’UTR (FarUTR), upstream of the start codon (PreStart), Start, ORF and Stop regions (Extended Data Table 1, schematized in Fig. 3a). As expected, we found that ribosome occupancy at multiple steps correlated well, indicating dependency in these cases (R^2^ higher than 0.74, Extended Data Fig. 4a, b and Supplementary Information). For instance, 30S subunits strongly correlated with 70S ribosomes at the Start region. T-30S_PreStart_ also correlated with T-30S_Start_, as did Free-30S and T-30S (both at PreStart or both at Start). We next asked whether certain SD motifs drive these correlations. For each correlated step, we analyzed SD sequences of mRNAs with the highest or lowest ratios between both states. This revealed a clear bias for stronger SDs in mRNAs with a high T-30S_PreStart_/Free-30S_PreStart_ ratio, much more pronounced than for any other comparison (Extended Data Fig. 4c, d). We also checked whether certain SD motifs were overrepresented in genes with high T-30S_PreStart_/Free-30S_PreStart_ ratios. We found that genes with the highest proportion of T-30S had a significant enrichment for an AGGAG SD sequence (Fig. 3b), even compared to other strong SD motifs (Extended Data Fig. 4c, Supplementary Information). Altogether, this suggests that the SD motif strongly influences 30S recruitment to the PreStart region and its assuming a primed state after the first translation cycle. This further suggests that this primed T-30S in the PreStart region is caused by preceding translating ribosomes.

**Figure 3.**
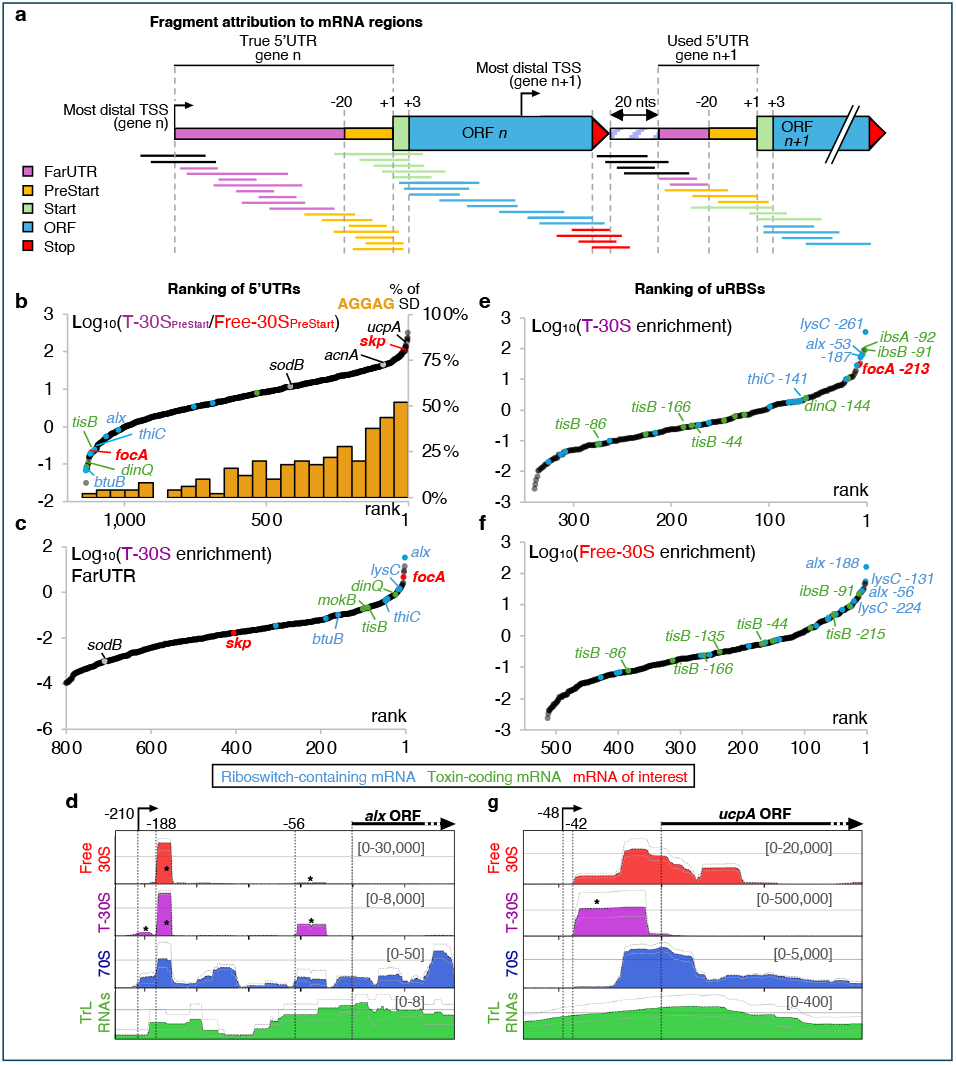
30S binds close and far upstream of TIRs within 5’UTRs. (**a**) Scheme of the different mRNA regions considered for the analysis of ribosome binding to mRNAs and criteria used for fragment attribution. (**b, c**) Ranking of mRNAs based on the highest T-30S_PreStart_/Free-30S_PreStart_ ratio (within positions - 20 to +2 relative to translation start, **b**) or the highest enrichment for T-30S in their FarUTRs (upstream of position -20, **c**). For T-30S_PreStart_/Free-30S_PreStart_ ratios, the percentage of mRNAs with an AGGAG SD motif are indicated by the orange histogram for each group of 50 genes. Are indicated here and in all subsequent rankings: toxin-coding mRNAs in green, riboswitch containing mRNAs in blue, and *focA* and *skp* in red as genes of interest. 30S-seq profiles of *alx* (**d**) and *ucpA* (**g**) mRNAs, as in Fig. 1d, e. **(e, f)** Ranking of uRBSs based on the highest T-30S (**e**) or Free-30S (**f**) enrichment. Note that the number of ribosome binding sites with 30S/70S ratios ≥ 5 is vastly enriched in FarUTRs (Extended data Fig. 7). Criteria used for selecting mRNAs (panels b, c) or binding sites (panels e, f) included in these analyses are indicated in Extended Data Table 6.

### The 30S subunit binds to hundreds of 5’UTRs at upstream Ribosome Binding Sites

We developed 30S-seq based on the hypothesis that 30S binding to 5’UTRs was underestimated. This was supported by the major enrichment of 5’UTR fragments in Free-30S and T-30S fractions (Fig. 1c). We therefore explored 30S binding to 5’UTRs in more detail. First, we ranked mRNAs based on their T-30S and Free-30S enrichment in FarUTRs (Fig. 3c and Extended Data Fig. 5a). Out of 916 mRNAs with measurable FarUTR T-30S levels, the known *E. coli* standby case *tisB* ranked 87^th^ (76^th^ among 801 mRNAs for the Free-30S). We thus consider that there is a very strong possibility that the 30S subunit binds to most higher ranking 5’UTRs and to at least some of the lower ranking 5’UTRs. Importantly, 30S enrichment only very poorly correlated with 5’UTR length (R^2^=0.387 and 0.294 for Free-30S and T-30S, respectively, Extended Data Fig. 5b, c). This excludes a passive 30S binding to free mRNA, and suggests an active and specific recruitment mechanism.

Strikingly, certain families of genes were over-represented in high ranking 5’UTRs. In particular, all known riboswitch-containing 5’UTRs as well as detectable toxin-coding mRNAs were among the top 20% T-30S- and/or Free-30S-enriched 5’UTRs (Fig. 3c and Extended Data Fig. 5a). Interestingly, mRNAs from these two families also showed distinct, specific 30S binding patterns. Indeed, toxin-coding mRNAs such as *tisB* (Fig. 1e) or *dinQ* (Extended Data Fig. 6a) displayed multiple 30S binding sites along their 5’UTRs. In turn, riboswitch-containing 5’UTRs such as the top ranking *alx* (Fig. 3d) or *thiC* (Extended Data Fig. 6b) most often harbored one or two major binding sites, far upstream of the translation start. Hence the 30S binding pattern to mRNA FarUTRs could be related to different modes of 30S recruitment or gene regulation. We refer to the different 30S binding sites in FarUTRs as upstream Ribosome Binding Site (uRBS) in the rest of the study.

Together, these data establish that multiple mRNA FarUTRs have one or more uRBSs. We next sought to thoroughly identify these uRBSs. For this, we first searched for Free-30S- or T-30S-covered regions of 15-45 nts in FarUTRs, PreStart and Start regions (Extended Data Fig. 7 and Table 2). Next, we filtered these data for 5’UTR binding sites (i.e., upstream of nt +3) that had 30S levels at least as high as the Start region of the same mRNA, and that had a ≥5-fold 30S/70S ratio (Fig. 3e, f and 4a, b, Extended Data Fig. 5d and 7 and Table 3).

When considering only the regions that are strictly within the FarUTR, this revealed 806 uRBSs, distributed in 497 different mRNAs, detected in at least one condition among the Free-30S or T-30S samples, with or without crosslink (Fig. 3e, f and 4a, b). In addition, multiple 30S binding-sites overlapped both the FarUTR and the PreStart regions. To ensure that they differed from the primed T-30S species observed in the metagene analysis (centered around the -20-5 region, Fig. 2b), we focused on overlapping sites that had a 5’end upstream of nt-35. We found 52 of them that we refer to as longRBSs as they appear to be contiguous with the canonical RBSs (Fig. 4a and Extended Data Fig. 5e, f). Overall, uRBSs or longRBSs are present in ∼25% of the 2189 genes with a known or potential 5’UTR of at least 35 nts (Fig. 4b). Messenger RNAs containing uRBSs or longRBSs were notably enriched in gene functions such as heat/temperature response and translation (Extended Data Fig. 6c). This could underly their specific mode of regulation or a need for strong gene expression. As already hinted by the general 30S enrichment in their FarUTRs, every known riboswitch-containing mRNA and most toxin-coding mRNA in *E. coli* carry uRBSs. In turn, several longRBSs were in mRNAs ranked among the 50 highest T-30S_PreStart_/Free-30S_PreStart_ ratios, i.e., *ucpA, acnA* and *skp* (Fig 3g, Extended Data Fig. 6d and see below for *skp*). This suggests a possible link between the longRBSs and a strong primed T-30S signal upstream of the start codon. Altogether, this uncovers a vastly underestimated breadth of 30S binding to bacterial mRNA 5’UTRs, both far and close upstream of TIRs.

**Figure 4.**
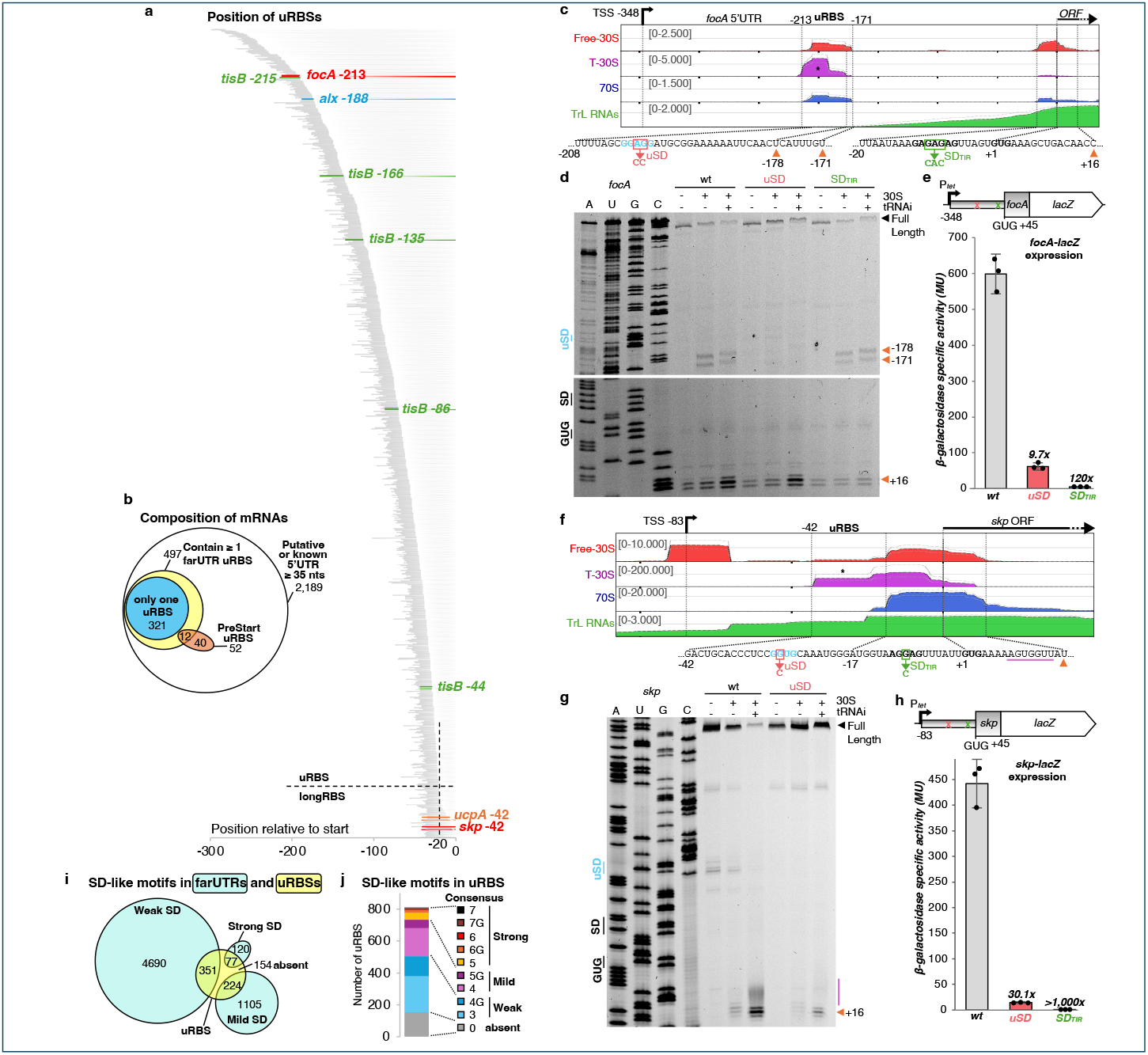
SD-like motifs within uRBSs and longRBSs affect 30S binding and gene expression. **a**. Position of uRBSs and long RBSs identified in this study relative to the translation start (0). The *focA* and *skp* uRBSs of interest are shown in red. 12 uRBSs found in long 5’UTRs upstream of position -300 are not displayed here. **b**. Venn diagram of uRBS-containing mRNAs among all mRNAs with known 5’UTRs of at least 35 nts or with a start codon separated by at least 55 nts from the closest upstream ORF stop codon. **(c, f)** 30S-seq profiles of *focA* (**c**) or *skp* (**f**) as in Fig. 1d, e. Sequences of the uRBS and of the TIR are depicted, with uSD motifs in blue and TIR SD (SD_TIR_) and start codons in bold. Mutations in the uSD motif or SD_TIR_ are depicted in red and green, respectively. The position of toeprinting signals obtained *in vitro* are represented by orange arrows for discrete sites and a purple line for more extended sites. **(d, g)** Toeprinting reactions on *focA* (**d**) or *skp* (**g**) were performed using *in vitro* transcribed mRNAs and cy5-labelled specific probes, in the presence or absence of 30S (20-fold excess) and/or tRNAi (40-fold excess). Sanger sequencing reactions are displayed on the left. **(e, h)** Beta-galactosidase assays were performed on strains carrying translational *P*_*tet*_*-focA-lacZ* (**e**) or *P*_*tet*_*-skp-lacZ* (**h**) gene fusions, wt or mutated in uSD or SD_TIR_. Repression fold-changes are indicated, and biological replicates are represented by a black dot. **i**. Venn diagram of the SD-like motif content of FarUTR uRBS and of the entirety of FarUTR regions analysed. Number of consensus nts in Weak SD: 3 nt or 4 nt with at least one A-to-G mismatch; Mild SD: 4 nt or 5 nt with at least one A-to-G mismatch; Strong SD: 5 to 7 nt, with or without A-to-G mismatch. The strong and mild SDs are, respectively, 5.2- and 2.2-fold overrepresented in uRBSs, with a hypergeometric P-value of 2.3×10^-36^ and 1.1×10^-33^. **j**. Distribution of SD-like motifs found in uRBSs. The number of consensus nts is indicated; G : the motif includes at least one A-to-G mismatch. Criteria used for selecting binding sites (panel a) or mRNAs (panels b, i and j) included in these analyses are indicated in Extended Data Table 6.

### 30S binding to uRBS or longRBS strongly impacts gene expression and often relies on SD-like motifs

To address the importance of 30S binding to uRBSs or longRBSs for gene expression, we selected a representative of each category for further study. For the uRBSs, we chose the *focA* mRNA, that encodes an inner membrane formate channel. Indeed, *focA* FarUTR was one of the most enriched for T- and Free-30S (Fig. 3c and Extended Data Fig. 5a). Furthermore, the *focA-213* uRBS (i.e., with a 5’end at nt-213) also ranked highly for T-30S enrichment (Fig. 3e), and was one of the most distant relative to the translation start (Fig. 4a, c and Extended Data Fig. 8a). We tested this site for 30S binding using *in vitro* toeprinting^9^ (Fig. 4d and Extended Data Fig. 8b). Remarkably, the 30S subunit bound to the same uRBS observed *in vivo* by 30S-seq, at a near-nt precision and independently of initiator tRNA (tRNAi). In contrast, 30S binding at the start region of *focA* required tRNAi. This is consistent with the absence of a functional start codon at the uRBS, and suggests that this binding can occur at a very early stage before the formation of a 30S initiation complex. It further suggests a very stable binding, following a mechanism at least partly different from binding at a TIR. We noticed that the uRBS contains a strong, conserved SD motif, hereafter referred to as upstream SD (uSD) (Fig. 4c and Extended Data Fig 8c). A mutation within this motif (*focA-uSD*) abolished 30S binding at the uRBS, with only a weak impact on binding at the TIR (Fig. 4d). Moreover, mutations in the weaker SD motif at the TIR (*focA-SD*_*TIR*_) prevented 30S binding at this region without affecting binding upstream, showing that 30S binding to both sites can occur independently. Finally, we tested the effect of these mutations on *focA* expression using a translational *focA-lacZ* fusion as a read-out (Fig. 4e). The expression of *focA* was affected ∼10-fold by the *focA-uSD* mutation, while the *focA-SD*_*TIR*_ mutation completely abolished gene expression. This demonstrates the major importance of both regions for *focA* expression.

As a representative of a longRBS-containing mRNA, we next chose to analyze *skp*, which encodes an essential periplasmic folding chaperone. *skp* stood out as having one of the highest T-30S_PreStart_/Free-30S_PreStart_ ratios (Fig. 3b) and displayed a longRBS starting at position -42, (Fig. 4a, f and Extended Data Fig. 5f and 8d). In agreement with the 30S-seq profile, toeprinting assays revealed a 30S-dependent smear upstream of the canonical initiation complex signal at position +16 (Fig. 4g). Similar to *focA*, we found a uSD motif within the *skp* uRBS, in this case 14 nts upstream of the TIR SD (Fig. 4f), and tested whether it could explain this 30S binding pattern. A point mutation in this motif (*skp*-*uSD*) led to an almost complete loss of both the smear profile and the +16 toeprint, indicating a strong involvement of the uSD motif in 30S binding (Fig. 4g). The same mutation led to a ∼30-fold repression of a *skp-lacZ* translational fusion *in vivo*, confirming its importance for *skp* expression (Fig. 4h). In turn, a point mutation in the TIR SD (*skp-SD*_*TIR*_) completely abolished *skp* expression, showing that the uSD does not function as a replacement SD but rather as an additional, major determinant of *skp* expression.

The importance of an SD-like motif in both *focA* and *skp* led us to search for similar motifs in all uRBSs and longRBSs. Of the 806 uRBSs identified, 80.9% (652) had a SD-like motif, with 43.5% (351), 27.8% (224) and 9.6% (77) showing a weak, mild or strong consensus sequence, respectively (Fig. 4i, j). In comparison with the global distribution of SD-motifs in FarUTRs, the strong and mild SDs were significantly overrepresented in uRBSs (Fig. 4i). In turn, we did not find a significant over-representation of SD-like motifs in longRBSs, indicating that they may only impact gene expression in rarer cases such as for *skp* (data not shown).

Hence, SD-like motifs are clearly a common feature of many uRBS and at least some longRBSs. As demonstrated for *focA* and *skp*, they likely play a crucial role for 30S recruitment and gene expression.

## Discussion

The 30S-seq approach described here uniquely establishes a map of the 30S binding sites in bacteria, both within and outside of the TIR, revealing new and unexpected insights into the translation initiation process. It presents various advantages over the previous transcriptome-wide translation initiation studies performed in bacteria. No antibiotic treatment was used for the 30S-seq, which avoids possible biases related to the antibiotic specificity. While the formaldehyde-crosslinking step is also susceptible to introduce biases, the dataset obtained in the absence of the crosslink step supports the same conclusions (Extended Data Fig. 7 and Extended Data Fig. 8a, d). In addition to the already major advances reported in this study, we now expect the 30S-seq approach to find many applications in the near future. In particular, it could become a method of choice to study translational control, to decipher the translation initiation process in detail in bacteria other than *E. coli*, in particular in those where translation is much less dependent on the SD sequence, or to address the importance of key factors in ribosome binding to different mRNA sites.

A first major advance brought by this study is the observation of multiple steps and conformations of 30S and 70S occupancy during initiation. This includes the translation of the first codons while the 70S is still attached to the mRNA SD, or the translation-dependent binding of the 30S subunit upstream of the start codon, primed to move into the initiation position. Interestingly, this primed 30S subunit is consistent with an individual gene study showing that the 30S subunit can be loaded to 5’UTR regions directly upstream of TIRs on the highly translated *lpp* mRNA^19^. In this case, after the first round of translation, subsequent 30S subunits can only access the TIR when entirely free of the leaving ribosome. The observation of a primed T-30S also perfectly corroborates a recent cryo-EM study demonstrating that the SD-antiSD interaction can take place at the exit of the mRNA tunnel, before an accommodation step flips the mRNA into the tunnel^20^ (Fig. 2c, step 2). This study was performed on a reconstituted translation initiation complex, in the presence of purified 30S subunits. However, according to our data, this mechanism is not restricted to the first translation cycle, and would in fact prevail in the second and later translation cycles. This mechanism would enable preparation for the next translation cycle as efficiently as possible.

Another major advance brought by the 30S-seq data is the identification of hundreds of uRBSs or longRBSs, that can play an important role in translation initiation (Fig. 3 and 4). Such non-canonical 30S binding sites are present in nearly 25% of the 2189 *E. coli* genes with at least 35 nts in their 5’UTR used for this analysis. This number is probably underestimated, as 30S signals in 5’UTRs overlapping with upstream ORFs were not considered, to exclude false positive signals coming from 70S dissociation within ORFs. Nevertheless, 30S binding sites in *bamA* 5’UTR, overlapping the *rseP* ORF (Extended Data Fig. 1g) or in *rpoS* 5’UTR, overlapping the *nlpD* ORF (Extended Data Fig. 6e), constitute examples of additional uRBSs to be explored.

Interestingly, our analysis identified uRBSs or longRBSs required for optimal expression in *focA* and *skp* mRNAs, both carrying a GUG start codon and other non-classical features in their TIR (see Supplementary Information). Other examples of uRBSs-carrying mRNAs, such as those containing riboswitches or encoding type 1 toxin mRNAs, are known to be highly structured mRNAs, in which the SD and the TIR may be poorly accessible. This could explain the need for uRBSs in these cases, with the structured regions predicted to be optimal for 30S binding. However, convincing uRBS profiles are not restricted to genes starting with non-conventional start codons (as *focA* or *skp*), or to a specific class of genes (riboswitches or toxins). Instead, they are found in hundreds of genes and are extremely diverse. Nevertheless, uRBSs or longRBSs appear enriched for several gene function categories, *e*.*g*., translation (Extended Data Fig. 6c), which could be related to specific mechanisms of gene regulation. Interestingly, this is clear for the ribosomal protein genes *rpsA* and *rpsO*: indeed, their uRBSs coincide with secondary structures or a pseudoknot in the mRNA known to impact 30S recruitment, respectively^11,21^ (Extended Data Fig. 6f, g). More generally, uRBSs provide an additional key region for bacterial gene expression and should form ideal targets for protein and RNA regulators, including riboswitches. This is already indicated by the extensive binding of the 30S subunit to riboswitch-containing mRNAs. It is also striking that several well-established examples of genes subject to translational control present one or more uRBSs in their 5’UTRs. This is true for genes controlled either by proteins, such as *thrS* or the afore-mentioned *rpsA* and *rpsO*, or by a large number of small RNAs, such as *rpoS* or *csgD*^22^ (Extended Data Fig. 6e, h).

Overall, the data presented here reveal that 30S binding in the 5’UTRs outside of the TIRS is far more frequent than previously anticipated. Such binding is reminiscent of the eukaryotic internal ribosome entry sites (IRES) that are structured mRNA regions within 5’UTRs recognized by the 40S and from which scanning can take place. From the 30S-seq data however, continuous footprints along 5’UTRs are visible only in very rare cases and over short distances. Thus, scanning along the 5’UTR from the 5’ end or from uRBSs seems unlikely or very rare, suggesting that the 30S could instead translocate from the 5’UTR to the TIR via binding to discrete sites. Future studies will be required to elucidate the mechanism of 30S loading, translocation and translation initiation for mRNAs carrying uRBSs. In other words, this study is the starting point to understand how the prokaryotic uRBSs compare to the role of 5’UTRs in eukaryotic translation.

## Methods

### Cells and media

*E. coli* strains used in this study are MG1655 (wt) or derivatives, as described in Supplementary Table 4. The different *lacZ* translational fusions were constructed by recombineering, as previously^23^, using primers listed in Supplementary Table 5. All cultures were performed at 37°C in LB, by inoculating fresh medium with a 1/500 dilution of overnight saturated cultures.

### Cell harvesting and lysis for 30S-seq

30S-Seq was performed following the framework described for yeast^4,5^, with significant adjustments inspired by RiboSeq in *E. coli*^24,25^ or set-up specifically in this study. For each of the three replicates, a 2 L culture of MG1655 strain reaching an optical density of 0.7 at 600nm was divided into two 1 L batches and poured onto 0.5 L of a frozen and crushed solution of 20 mM Mg-Acetate, under strong agitation. One batch was immediately crosslinked with the addition of formaldehyde (Thermo Scientific 28908) to a final concentration of 0.25%. After 10 min incubation with agitation, cells were pelleted down by centrifugation for 4 min, and the supernatant discarded. Exactly 20 min after initial formaldehyde addition, pellets were resuspended into ice-cold buffer A (25 mM Hepes-KOH pH 7.6, 100 mM NH_4_-Acetate, 20 mM Mg-Acetate, 2 mM β-Mercaptoethanol) supplemented with 25 mM Glycine to quench the remaining formaldehyde. At ice-cold temperature, cells were further washed and pelleted down twice in A buffer, then resuspended in 600 μL of buffer A. 150 μL of lysozyme at 20 mg/mL was added to lyse the cells for 2 min, followed by 30 μL of DNase I (Qiagen 79254) at 1 μg/μL, 90 μL of Brij 10% detergent and 90 μL of deoxycholic acid 10% to avoid ribosome aggregation with chromatin and to increase polysome yields. Lysates were centrifugated twice to eliminate cell debris, and total RNA concentration was estimated in cleared lysates by measuring its OD_260_.

### Ribosome sedimentation and MNase digestion

Translated RNAs were isolated by ultra-centrifugation of 3 mg total RNAs on 5%-15% sucrose gradients, buffer A, in 4 mL quick-seal tubes (Beckman Coulter 358650) for 65 min at 41 krpm and 0°C in a SW 41 Ti Swinging-bucket rotor, averaging 207,219 g. Pellets were recovered in 1 mL buffer A supplemented with 5 mM CaCl_2_ and the translated RNA concentration was determined. 400 to 500 μg of translated RNAs were then digested for 1h at 25°C and 1400 rpm shaking with 0.075 U of Micrococcal S7 Nuclease per μg of RNA (MNase, concentration of 1,9 U/μL) (Roche 10107921001) supplemented with 0.24 U of SUPERase-In RNase Inhibitor (Invitrogen AM2696) per μg of RNA. Digestion reactions were then stopped by quenching CaCl2 with 5 mM EGTA pH 8.0. T-30S and 70S were isolated by fraction collection after ultra-centrifugation of MNase-digested translated RNAs on 7.5%-30% sucrose gradients, buffer A, in 12 mL open-top tubes (Seton Scientific 7030) for 14h45 at 20 krpm, 0°C in a SW 41 Ti Swinging-bucket rotor, averaging 49,304 g. Similar gradients were used to separate Free-30S fractions from 500 μg of total RNAs. These fractions were then MNase-digested as described above, but in conditions adjusted to a ∼50 fold more dilute sample, with 15 U MNase per μg of RNA and a concentration of 0,1 U/μL.

### Reverse-crosslinking and mRNA protected fragments purification

Free-30S, T30S and 70S fractions as well as 20 μg of translated RNAs and 50 μg of total RNAs were reverse-crosslinked for 45 min at 65°C, 1300 rpm shaking, in a buffer containing 1% SDS, 10 mM EDTA pH 8.0, 10 mM glycine and 10 mM Tris HCl pH 7.4, and 1 volume of phenol-chloroform. Samples were then centrifuged for 5 min at 13.2 krpm, and the aqueous phase was recovered. 70S, translated and total RNA samples were phenol-extracted again, and all samples were next precipitated with 2 μL glycoblue (Invitrogen AM9515) and 1.3 volumes of isopropanol. RNA pellets were ethanol washed, resuspended in water, and quantified by measuring the absorbance at 260 nm. 1 μg of Free-30S-, T-30S- or 70S-associated RNA fragments were run on 10% polyacrylamide denaturing gels, and 15 to 45 nt-long ribosome footprints were gel-purified.

### RNA preparation for sequencing (ribosome profiling and total RNAseq)

Ribosome-associated footprints were denatured for 2 min at 80°C, 3’-dephosphorylated for 1 h at 37°C with 10 U (1 U/μL) of T4 PNK (New England Biolabs M0201), and 5’-phosphorylated by adding 1 mM ATP to the reactions and incubating for another 30 min. Libraries construction and sequencing were performed by the iGenSeq core facility, at the ICM (Institut du Cerveau et de la Moelle Epinière, Paris). Ribosome profiling samples were prepared with the QIAseq miRNA 96 Index Kit IL UDI-A (Qiagen 331905), without fragmentation, but with an extra step of ribodepletion using the QIAseq FastSelect -5S/16S/23S kit (Qiagen 335925), and sequenced on a NovaSeq 6000 (Illumina), set for 100 nt-long paired end reads. Input RNAseq was performed using 1 μg of total RNAs or translated RNAs using the QIAseq Stranded RNA Lib Kit UDI (Qiagen 180450), with an extra step of ribodepletion using the QIAseq FastSelect -5S/16S/23S kit (Qiagen 335925), and sequenced on a NextSeq 2000 (Illumina), set for 50 nt-long paired end reads.

### Sequencing analysis

Adapter sequences were trimmed off from the miRNA fraction using the cutadapt tool v4.2^26^. Additional hard-clipping was performed on r2 in order to remove degenerated sub-sequences from the QIAseq miRNA 3’ adapter. Read pairs with an average sequencing quality under 30 for r1 and 28 for r2 were discarded. The trimmed miRNAs mapping to the seven ribosomal operons of *E. coli* NC_000913.3 were filtered out. Other reads were then realigned on the complete genome using bowtie2 v2.5.0^27^ (−-maxins 400 -d 15 -r 3 -n 0 -l 15 -i S,1,0.50). Reads below 13 bp were removed to avoid potential mapping artifacts. The Total and Translated mRNAs were mapped with bowtie2 v2.5.0 (−-maxins 2000 --local –very-sensitive-local) on *E. coli* NC_000913.3.

### 30S-seq quantification

Ribosome-associated RNA fragments feature quantification was performed either on uniquely- and multi-mapped reads for rRNAs, tRNAs and SsrA, or on uniquely-mapped reads (mapQ >= 20) for mRNAs and remaining ncRNAs. For uniquely mapped reads, quantification results were categorized into the following regions: Start (reads contain the entire first codon of the ORF), Stop (reads contain the entire last codon of the ORF), ORF (reads aligned anywhere only within the ORF and is not Stop or Start), FarUTR (reads aligned anywhere from the TSS position to the -20 position) and PreStart (3’end of reads between position -19 and +2) Total and Translated mRNAs feature quantification was performed using featureCounts v2.0.3^28^. GFF, TSS coordinates and operons were obtained from ecocyc^29^. Conditions for mRNA incorporation into analysis are detailed in Supplementary Table 6.

### Peak detection

In order to detect unconventional 30S binding sites in 5’UTRs, the pileup of the miRNA fraction (obtained by using the samtools mpileup tool v1.16.1^30^ was processed by the find_peaks function from the scipy.signal module v1.14.1. For each 5’UTR, peak detection was set to at least 10% of the read count at the start codon. Common peaks between samples were sorted into two categories called “FarUTR” and “PreStart”, with 3’ ends respectively before or at, and after position -20 relative to translation start. Conditions for ribosome binding sites incorporation into analysis are detailed in Supplementary Table 6.

### Toeprinting Assays

Toeprinting assays were performed essentially as described previously^13^. Briefly, RNAs were prepared by *in vitro* transcription with a T7 polymerase (Invitrogen AM1334) using a DNA template generated by PCR to introduce a T7 promoter (primers sequences in Supplementary Table 5). 0.5 pmol of RNA and 2 pmol of Cy5-labelled probe were denatured at 80°C for 3 min, flash-frozen in solid CO_2_/ ethanol for 1 min, then brought to 1x AMV buffer (75 mM K-Acetate, 50 mM Tris-HCl pH 8.3, 8 mM Mg-Acetate, 10 mM DTT) and annealed at 37°C for 5 min. After supplementation with 15 nM dNTPs, and with 15 pmol tRNA^fmet^ and 10 pmol 30S ribosomal subunits whenever indicated, the reactions were further incubated at 37°C for 10 min. Finally, reverse transcription was carried out using 0.1U/μL of AMV reserve-transcriptase (New England Biolabs M0277) in the reactions for 20 min at 37°C, before reactions were stopped by the addition of a formamide/EDTA mixture. cDNAs were observed after migration on a 6% denaturing PAGE, using a Typhoon fluorescent scanner set for Cy5 detection.

### β-galactosidase Assays

β-galactosidase assays were performed on cells grown to an optical density of 0.4 at 600nm, as described previously^31,32^, using chloroform to lyse the cells. The β-galactosidase specific activity was calculated using the Miller formula. Data are shown as average of three independent replicates, and error bars indicate intervals of confidence.

## Acknowledgements

We thank Ciarán Condon and Mathias Springer for careful reading and insightful comments on the manuscript. We are grateful to members of the team *RNA control of gene expression* for discussions and support. We thank Claude Chiaruttini for initial help with the 30S purification procedure, and Gregory Boël and members of his group for initial help with the sucrose gradients collection. This work benefited from equipment and services from the iGenSeq core facility, at ICM. This project has received funding from the European Research Council (ERC) under the European Union’s Horizon 2020 research and innovation program [grant agreement no. 818750] and under the Marie Skłodowska-Curie grant agreement No 101034407; research in the UMR8261 is also supported by the CNRS and the ‘Initiative d’Excellence’ program from the French State [‘DYNAMO’, ANR-11-LABX-0011].

## Author contributions

JJ and MG designed the study and supervised the work, JJ developed and performed the bacterial 30S-seq experiment, VGN and AM developed and performed the bioinformatical analysis of the 30S-seq, MAS and JJ performed toeprinting and β-galactosidase experiments, JJ and MG wrote the first draft, and all authors discussed results, reviewed, and edited the manuscript.

