## Supplementary information for "30S-seq redefines the bacterial Ribosome Binding Site"

**Supporting text for Fig. 1 and Extended Data Fig. 1 and 2:**
**30S-seq successfully identifies 30S- and 70S-binding sites in *E. coli***

The quality of the 30S-seq results was assessed with multiple indicators. First, we verified the reproducibility between replicates using a primary component analysis (PCA) on mRNA fragments found in each sample (Fig. 1b). We next checked fragments distributions (Fig. 1c, Extended Data Fig. 1b) and sizes (Extended Data Fig. 2) in the Free-30S, T-30S and 70S fractions. They clearly differed in the three different fractions, with most changes between Free-30S and 70S, while the T-30S fractions showed more similarity with Free-30S. As expected, the ratio of fragments from small subunit rRNA (16S) to large-subunit rRNAs (23S+5S) was higher in the 30S fractions (~4.5 and ~0,2 in Free-30S and T-30S samples, respectively) compared to the 70S (less than 0,02) (Extended Data Fig. 1b). The relatively low T-30S ratio (compared with Free-30S) could be due to early 70S assembly steps found in T-30S fractions, as well as contacts with the subsequent elongating 70S and/or dissociation of unstable elongating or terminating 70S. In these conditions, the 23S rRNA could contact the 30S subunits and be vulnerable to MNase digestion, leading to its recovery with T-30S particles. The rRNA leader sequences provided another indication of the quality of the different fractions. Indeed, we expected Free-30S fractions to include non-fully mature particles, possibly unbound to mRNA. This is visible by 16S transcript leader fragments that are ~37.5-fold and >1000-fold higher than in the T-30S and 70S fractions, respectively (Extended Data Fig. 1f).

In turn, tRNA fragments represented ~45% of total fragments in Free-30S fractions and more than 20% of fragments in T-30S, vs less than 2% in 70S fractions (Fig. 1c). The large number of rRNA reads in the 70S fraction possibly contributes to the apparent decrease of the other RNAs, including the tRNAs. In addition, this low proportion of tRNAs in the 70S samples is also likely due to a different interaction of the tRNAs with the 70S particles than with the 30S alone, resulting in a different MNase digestion pattern. This is further shown by the size of the tRNA fragments in the 30S-seq analysis: there is a majority of 31-33 nts-long fragments in Free-30S and T-30S fractions vs 24-26 nts-long in 70S fractions (Extended Data Fig. 2).

When looking at the distribution of the protected fragments on the mRNAs, 70S were present almost exclusively on the coding sequences (protecting from the TIR to the stop codon). In contrast, proportions of mRNA 5'UTR fragments in Free-30S and T-30S fractions were much higher than in 70S fractions: ~12% and up to 14.5% in Free-30S and T-30S fractions, respectively, vs less than 0.5% in 70S fractions (Fig. 1c). This shows that 30S frequently binds 5'UTRs. It is furthermore important to highlight that reads mapping to 5'UTRs overlapping with ORFs were counted as ORFs. As a result, the number of 5'UTR fragments is under-estimated, while that of ORFs fragments is over-estimated. This is expected to be especially important in the Free-30S and T-30S fractions, but not so much for the 70S given the very poor coverage of 5'UTR in this fraction. For instance, the 902 nt-long *bamA* 5'UTR extensively overlaps with the preceding *rseP* ORF, and is almost exclusively bound by 30S particles. (Extended Data Fig. 1g) The corresponding reads yet map to ORF fragments rather than 5'UTRs, even though the translation in this region is very low, as indicated by the low number of 70S reads (Extended Data Fig. 1c).

**Supporting text for Fig. 3 and Extended Data Fig. 4 :**

**Strong SD motifs correlate with 30S subunits primed upstream of the start codon**

We analysed the correlation between the enrichments of the different fractions (Free-30S, T-30S or 70S) in the Prestart and Start regions (Extended Data Fig. 4a). We found that 70S enrichment correlated well with both Free-30S$_{Start}$ ($R^2$=0.841) and T-30S$_{Start}$ ($R^2$=0.887). This indicates as expected that efficient 70S assembly depends on 30S accommodation to the start site. Free-30S$_{Start}$ and T-30S$_{Start}$ also correlated strongly ($R^2$=0.893), indicating that most mRNAs accommodate well 30S at both the first and subsequent translation cycles. Furthermore, T-30S$_{Start}$ correlated well with T-30S$_{PreStart}$ ($R^2$=0.743). This suggests that T-30S enrichment at PreStart is a good indicator of T-30S accommodation to the Start. Therefore, primed T-30S upstream of the start codon could increase the accommodation rate, and hence 70S assembly and translation initiation. In turn, Free-30S$_{Start}$ and Free-30S$_{PreStart}$ correlated poorly ($R^2$=0.612), showing that this mechanism likely depends on the presence of a preceding 70S.

Surprisingly, Free-30S$_{PreStart}$ and T-30S$_{PreStart}$ also correlated well ($R^2$=0.810). In addition, Free-30S$_{PreStart}$ levels were generally ~0.6 log lower than T-30S$_{PreStart}$ (Extended Data Fig. 4b), indicating that Free-30S subunits frequently cover PreStarts a fraction of the time the T-30S subunits do. We reasoned that the PreStart coverage time of Free-30S vs T-30S could be influenced by certain SD motifs, and compared mRNAs by their ratio of T-30S$_{PreStart}$/Free-30S$_{PreStart}$ (Fig. 3b). We found that mRNAs with the highest proportion of T-30S had a significant enrichment for the AGGAG SD (Fig. 3b and Extended Data Fig. 4c). Indeed, out of the 50 mRNAs with the highest ratio, 26 had an AGGAG motif, representing a 3.43-fold enrichment with a probability of occurring at random of $4.59\times10^{-10}$. In turn, only one mRNA within the 50 lowest ratios of T-30S$_{PreStart}$/Free-30S$_{PreStart}$ had an AGGAG motif : *tisB*, which had cryptic levels of 70S within its ORF, likely due to its constitutive repression[1,2]. Interestingly, most mRNAs within the 50 lowest ratios had ill-defined SD motifs (Extended Data Fig. 4d). No other group of mRNAs had a SD motif-bias as dramatic as that of high T-30S$_{PreStart}$/Free-30S$_{PreStart}$. However, we noticed that mRNAs with lowest 70S$_{Start}$/T-30S$_{Start}$ had stronger SD motifs that those with a high ratio (Extended Data Fig. 4d). Altogether, this suggests that the strength as well as the specific sequence of the SD directly impacts the occupancy of primed 30S at PreStart after the first round of translation, most likely via the presence of preceding translating ribosomes. In turn, stronger SD appear detrimental for rapid 70S assembly, in agreement with previous reports[3,4].

**Supporting text for Fig. 4:**

**Specificities of the *focA* and *skp* mRNAs and possible relation with their uRBS or longRBS**

Our studies strongly indicate that the uRBS or longRBS and the uSD motifs such as those found in *focA* and *skp* help to increase 30S recruitment efficiency. This could be especially important when the TIR is already occupied by an initiating ribosome. As both genes start with a GUG start codon, it is possible to imagine that initiation takes longer on these mRNAs. *skp* also possesses a strong AGGAG SD motif, showing the typical features of an mRNA with a very high T-30S$_{PreStart}$/Free-30S$_{PreStart}$ ratio (Fig. 3b). In this case, the requirement for 30S binding upstream of the TIR could be related to the strong occupancy of T-30S at the PreStart region, possibly limiting the direct recruitment of 30S to the TIR. The *focA* TIR is different, with three overlapping poor GAG SD motifs and two overlapping non-canonical GUG start codons (Fig. 4c and Extended Data Fig. 8c). The first GUG starts at position -2, but out-of-frame translation at this position would be immediately aborted by the following UGA stop codon. This may lead to inefficient translation initiation. Hence, both *focA* and *skp* translation may require an alternative pathway for 30S recruitment that does not rely solely on the TIR. Future studies, not restricted to *focA* and *skp*, should reveal how 30S binding to uRBSs or longRBSs differs from binding to TIRs– such as, for instance, the observed tRNAi-independent binding to uRBS (Fig. 4d and Extended Data Fig. 8b)– and how this participates in translation.

### Supplementary Figures

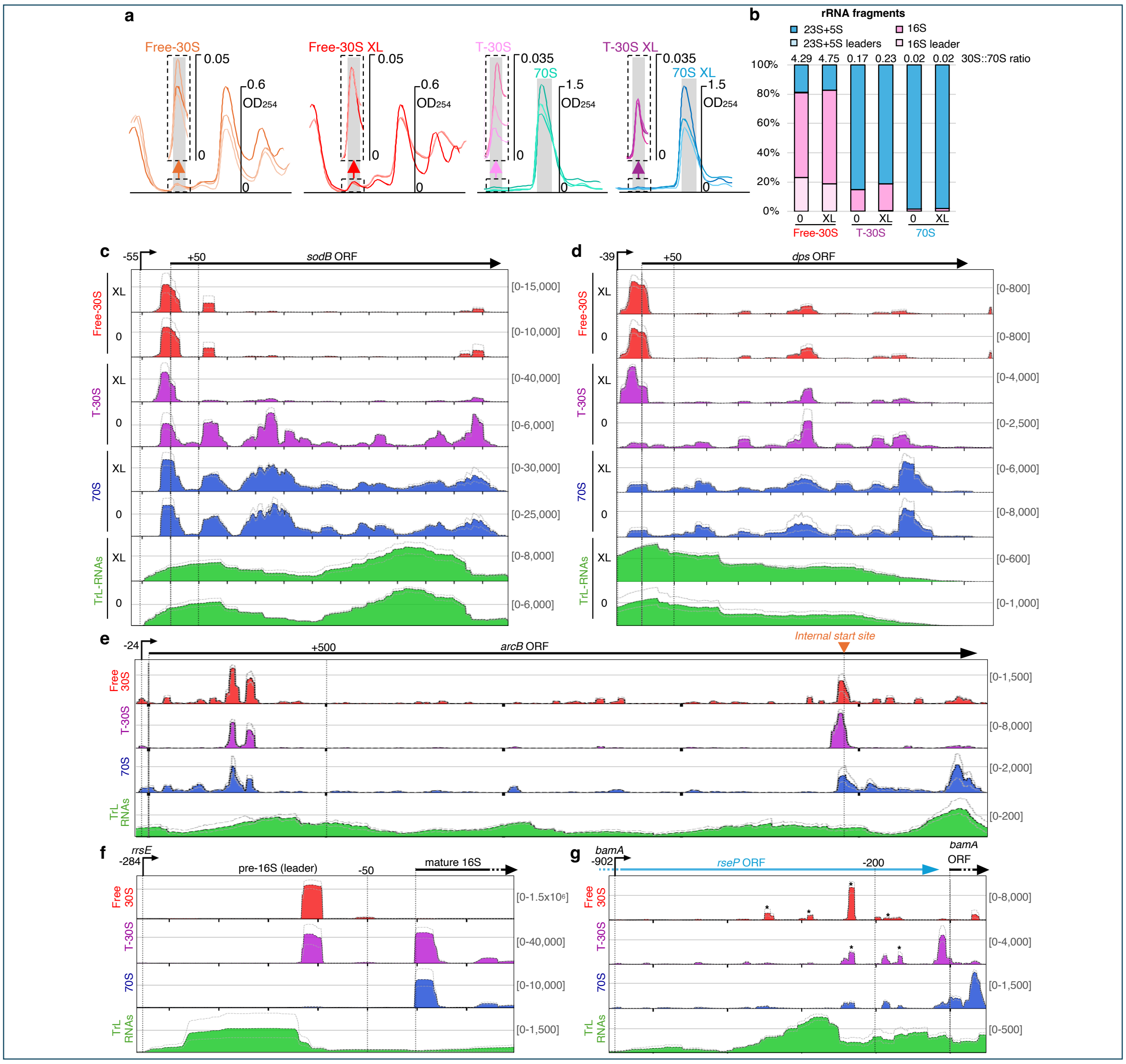


**Extended Data Figure 1 | Establishment of the 30S-seq. a.** Sedimentation gradient profiles of different fractions separated for the 30S-seq. Biological triplicates are shown stacked, and scales (optical density at 260 nm) are displayed for the top sedimentation profile. **b.** Repartition of protected fragments aligning to rRNAs in Free-30S, T-30S or 70S fractions in crosslinked (XL) or non-crosslinked (0) samples. The ratio of (16S)/(23S + 5S), including the leader and mature sequences, is represented (30S::70S ratio). **(c-g)** 30S-seq profiles of *sodB* (**c**), *dps* (**d**), *arcB* (**e**), 16S (*rrsE,* **f**) and *bamA* (**g**), as in Fig. 1d, e. For *sodB* and *dps*, crosslinked (XL) and non-crosslinked (0) samples are displayed. For *arcB*, the known internal start site[5] is indicated by an orange arrow. For *rrsE*, the 16S leader and mature regions are shown instead of 5'UTR and ORF; the observed profile is the leftover result of sequencing after rRNA removal (using the QIAseq FastSelect -5S/16S/23S kit). For *bamA*, the preceding ORF (*rseP*) is displayed in blue. *: sites that would be considered uRBSs (see description below) if they did not overlap the preceding ORF.

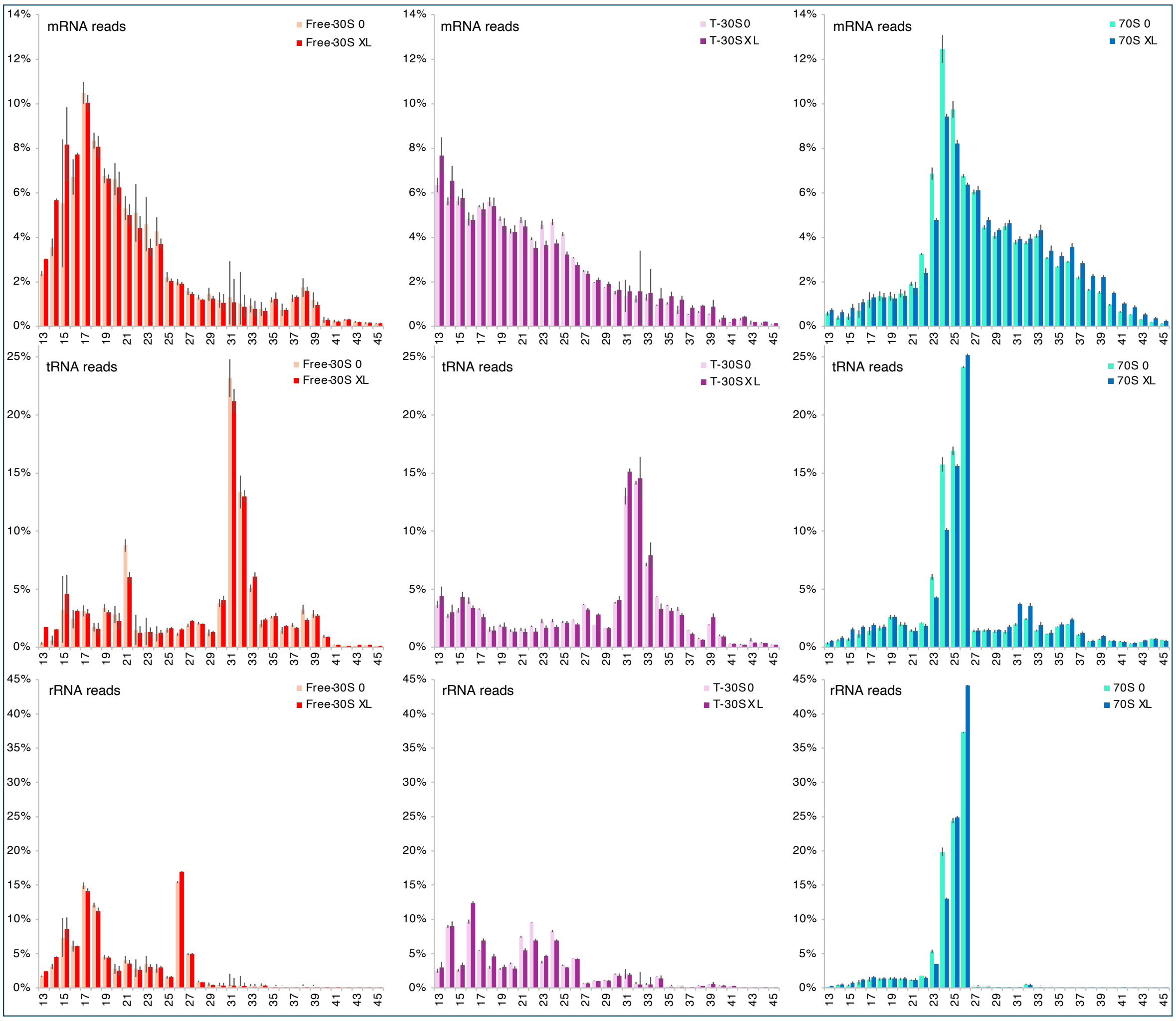


**Extended Data Figure 2 | 30S and 70S protections generate different fragment sizes.** Distribution of protected RNA sizes in crosslinked (XL) and non-crosslinked (0) samples of the Free-30S, T-30S and 70S fractions.

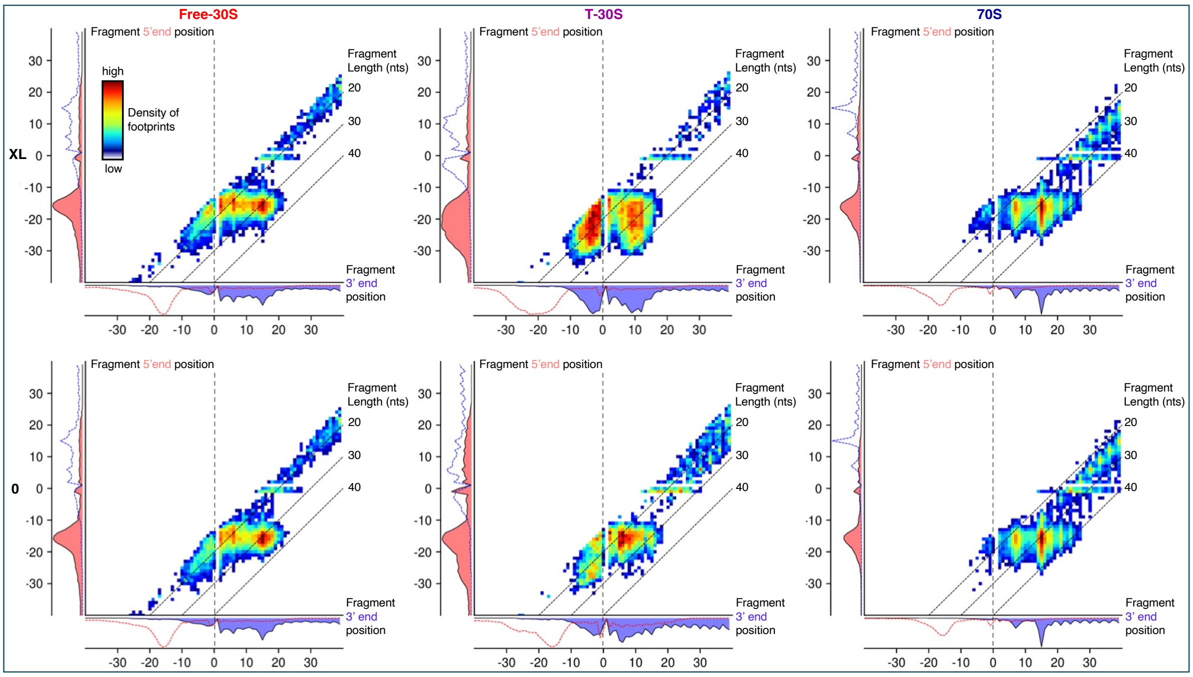


**Extended Data Figure 3 | Crosslinking stabilizes T-30S at PreStart.** Heatmap of Free-30S, T-30S and 70S protected fragments extremities in a subset of 1919 genes that are first in their operons with at least 15 fragments at the start region in at least one of the Free-30S, T-30S or 70S fractions, as in Fig. 2a-c. Results for crosslinked (XL, top panels, also displayed in Fig. 2) and non-crosslinked (0, bottom panels) samples are displayed. For non-crosslinked samples, 1848 genes were considered based on the same criteria of at least 15 fragments at the start region in at least one of the fractions.

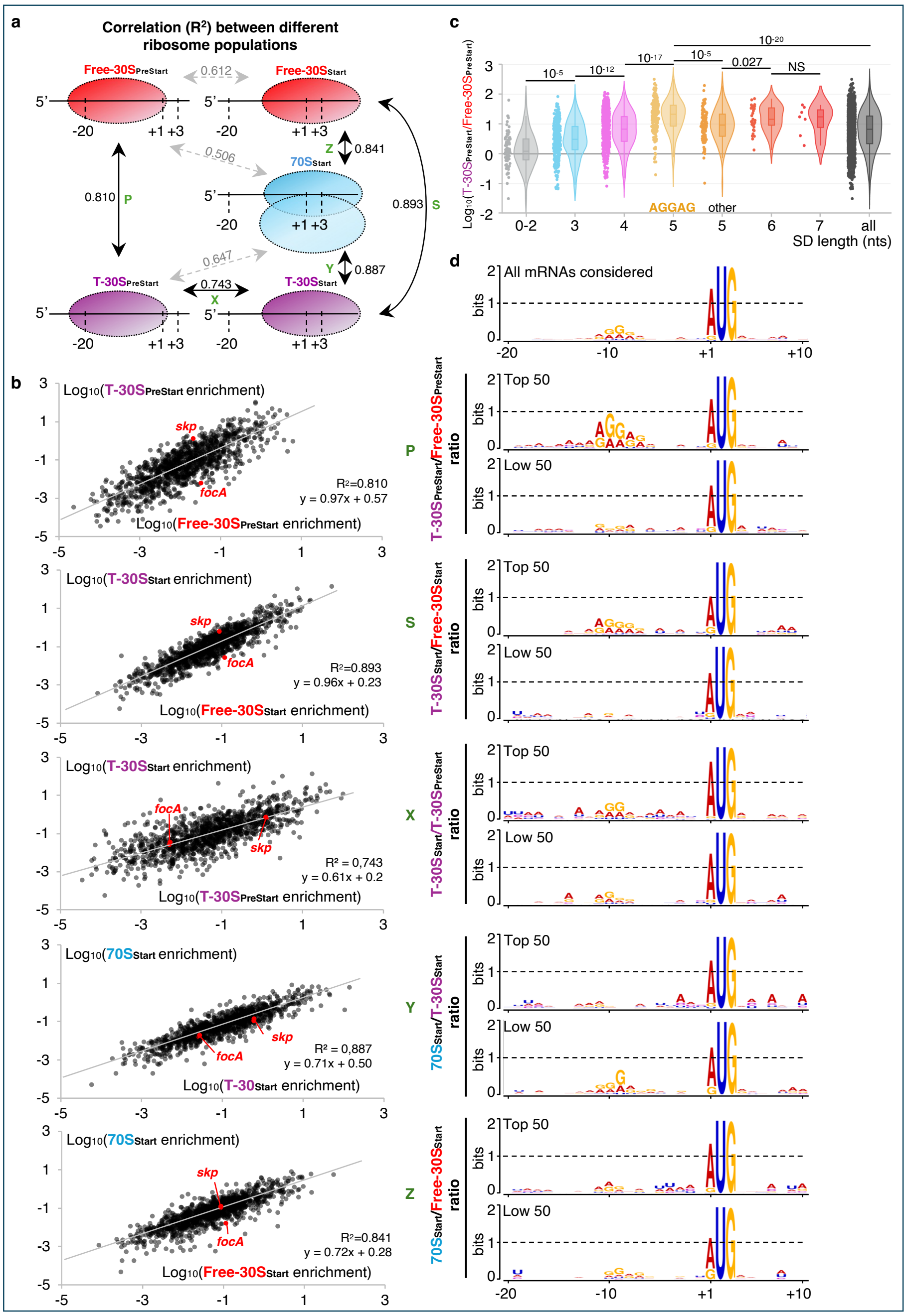

**Extended Data Figure 4 | Relationship between ribosome populations at TIRs. a.** Representation of the Pearson ($R^2$) correlation between the enrichment of different 30S samples at PreStart and Start, or 70S at Start. Black arrows represent the strongest (>0.7) correlations. P: T-$30S_{PreStart}$ vs Free-$30S_{PreStart}$. S: T-$30S_{Start}$ vs Free-$30S_{Start}$. X: T-$30S_{Start}$ vs T-$30S_{PreStart}$. Y: $70S_{Start}$ vs T-$30S_{Start}$. Z: $70S_{Start}$ vs Free-$30S_{Start}$. **b**. Pair-wise (P, S, X, Y, Z) scatter plots of the indicated sample enrichments. Trend curves are displayed, as well as the corresponding equation. **c**. Comparison of the T-$30S_{PreStart}$/Free-$30S_{PreStart}$ ratio in genes with different SD motifs shown as a violin plot (https://datatab.net/statistics-calculator/charts/violin-plot). The P-value statistical significance between mRNAs with different motifs was estimated using bilateral homoscedastic Student T-tests and is indicated on top. **d**. TIR (-20 to + 10 relative to translation start) sequence WebLogo analysis (https://weblogo.berkeley.edu/logo.cgi[6]) of the 50 mRNAs with the highest or lowest indicated ratios (P, S, X, Y, Z). Criteria used for selecting mRNAs included in these analyses are indicated in Extended Data Table 6.

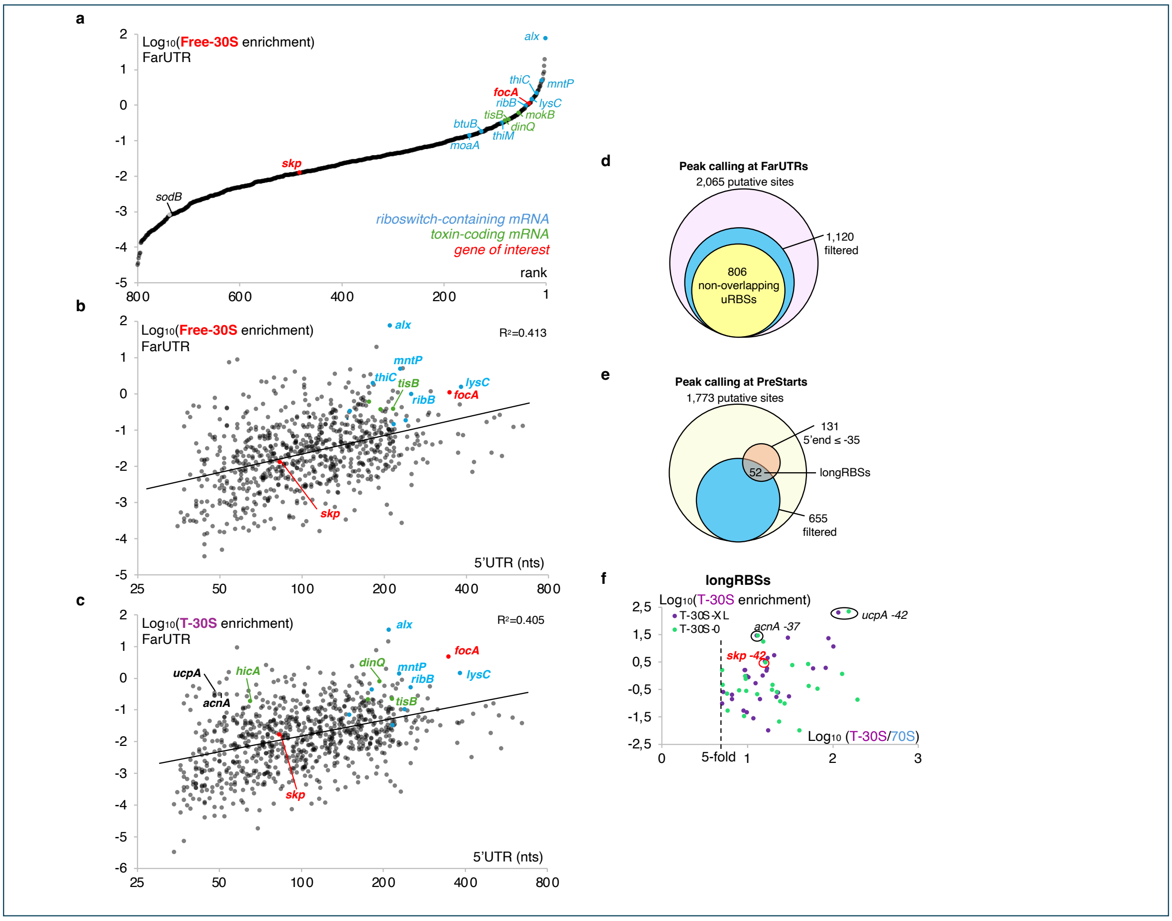


**Extended Data Figure 5 | Quantitation and position of 30S binding to 5'UTRs.** Ranking of mRNAs based on the highest enrichment for Free-30S in their FarUTRs (upstream of position -20), as in Fig. 3c. Scatter plots of Free-30S (**b**) or T-30S (**c**) FarUTRs enrichment vs 5'UTR length. The $R^2$ Pearson correlation and trend curves are displayed. Riboswitch-containing mRNAs are in blue, toxin-coding mRNAs in green and the *focA* and *skp* genes of interest in red. Venn diagrams represent the total number of peaks that qualify as uRBS (**d**) or longRBSs (**e**), according to the conditions described in the text. **f**. Scatter plot of crosslinked (XL) and non-crosslinked (0) T-30S enrichment vs T-30S/70S ratios in longRBSs. Criteria used for selecting mRNAs or binding sites included in these analyses are indicated in Extended Data Table 6.

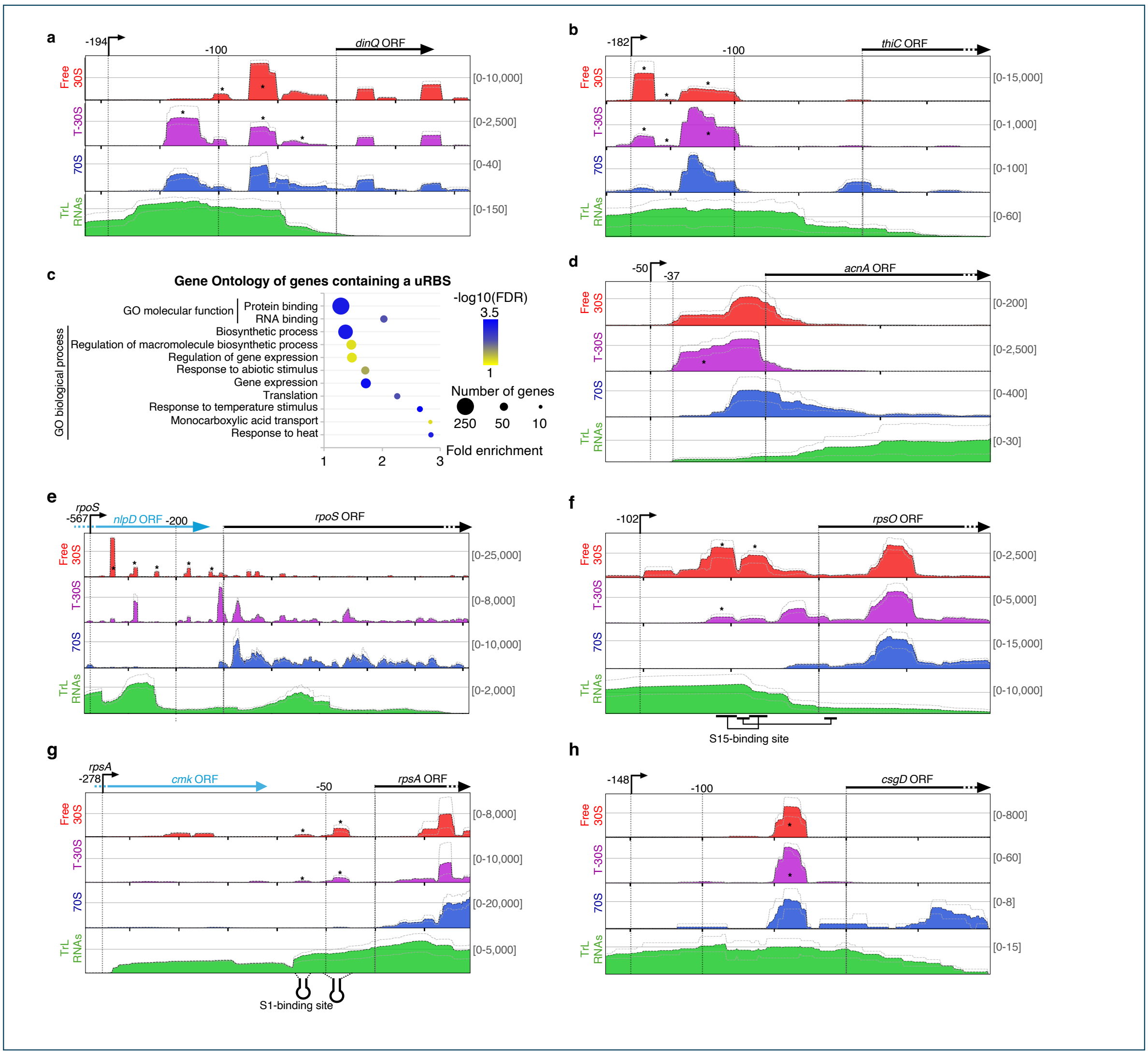


**Extended Data Figure 6 | Diversity of profiles and mRNAs containing uRBSs.** 30S-seq profiles of *dinQ* (**a**), *thiC* (**b**), *acnA* (**d**), *rpsA* (**e**), *rpsO* (**f**), *csgD* (**g**) *and rpoS* (**h**), as in Fig. 1d, e. For *rpsA* and *rpoS,* the preceding ORFs (*cmk* and *nlpD,* respectively) are displayed in blue. For *rpsA* and *rpsO*, known specific S1- and S15-binding sites are shown[7,8]. *: sites detected as uRBSs (see description below) or that would be considered uRBSs for *rpoS*, if they did not overlap with the preceding ORF. **c**. Gene Ontology analysis of mRNAs containing uRBS(s), according to the criteria described in Extended Data Table 6. FDR: False Discovery Rate.

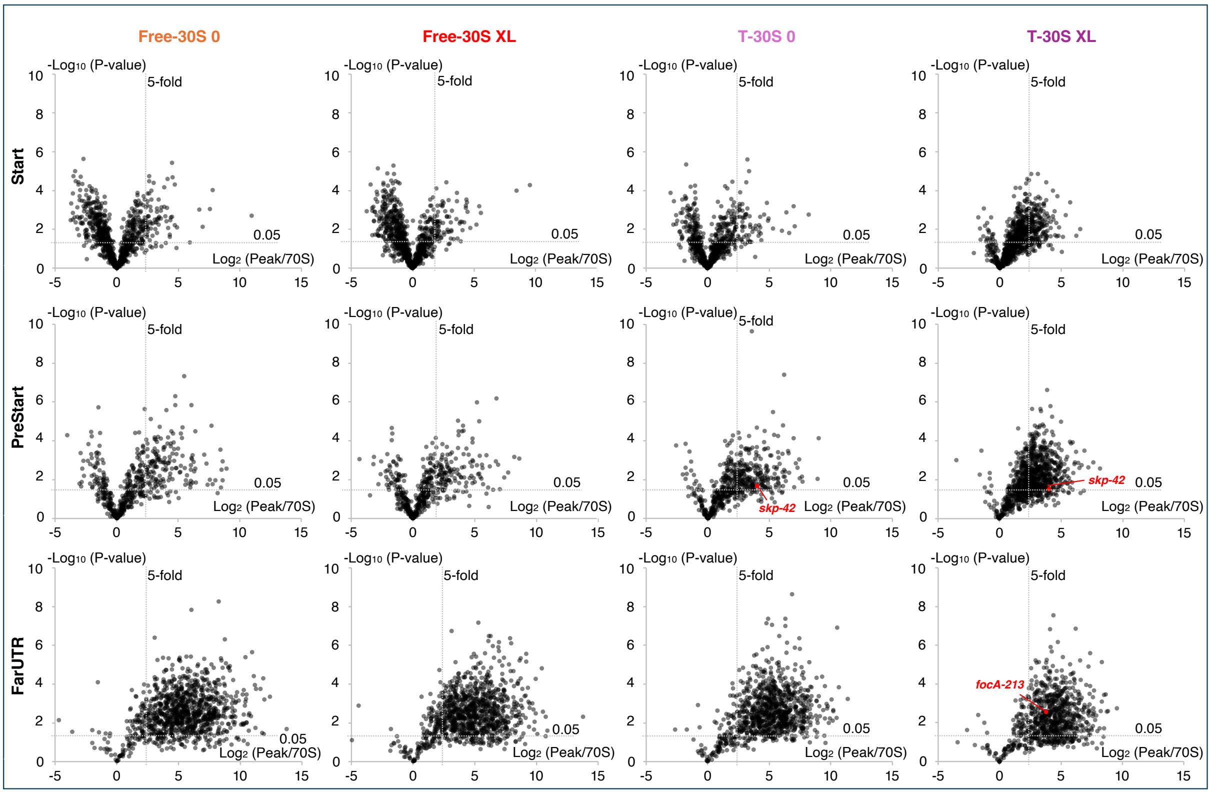


**Extended Data Figure 7 | uRBS are poorly covered by 70S.** Volcano plots of the 30S *vs* 70S signals at detected peak in the indicated sample and mRNA region. The ratio (5-fold) and P-value (0.05) thresholds used to consider a peak in FarUTR or at PreStart as a uRBS are displayed as dotted lines.

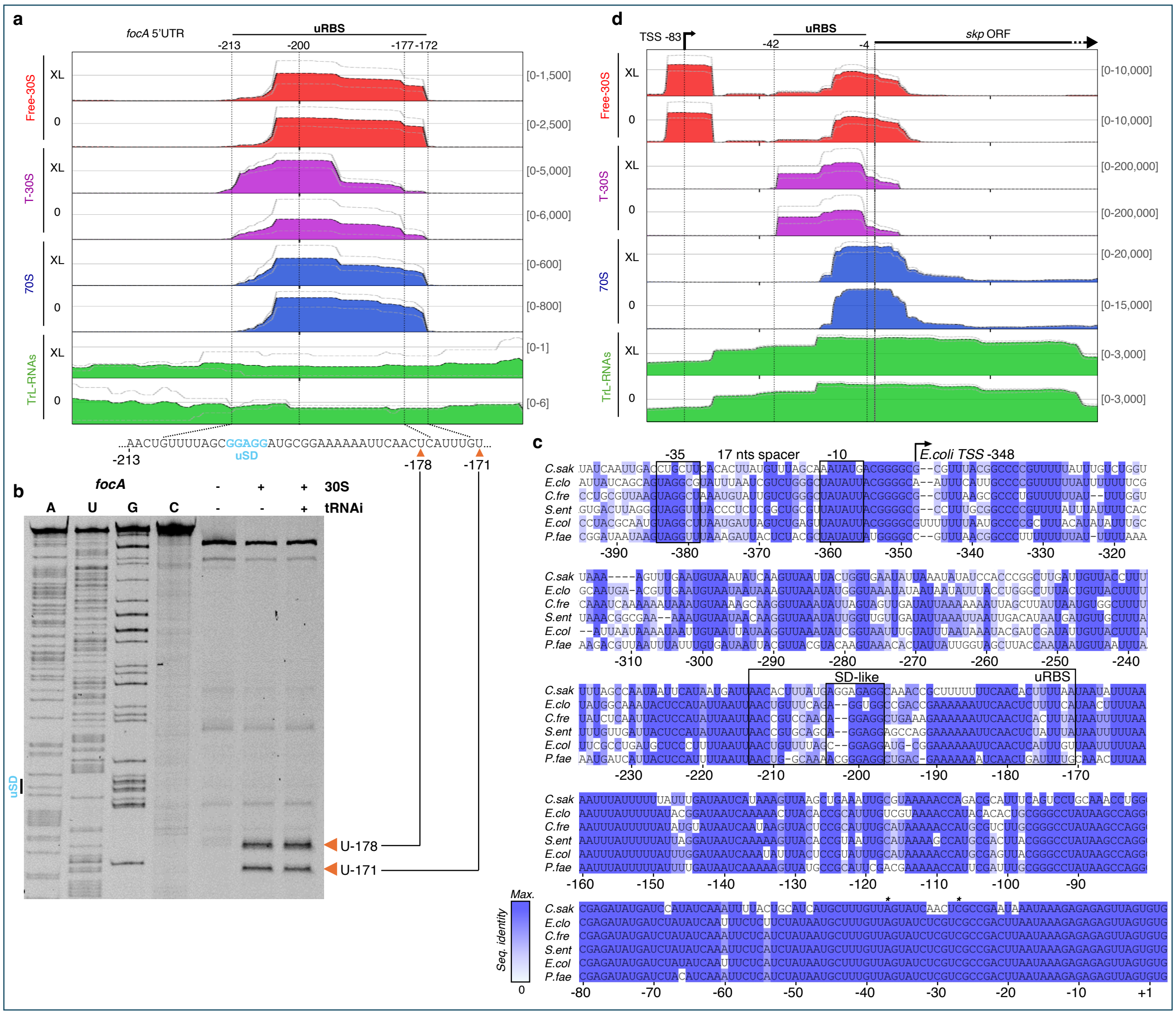


**Extended Data Figure 8 | Crosslink dependence, precision and conservation of 30S binding to uRBSs.** 30S-Seq profiles of *focA* (**a**) and *skp* (**d**), as in Fig. 1d, e. Crosslinked (XL) and non-crosslinked (0) samples are displayed. **b**. Toeprinting assays on *focA,* as in Fig. 4b. The Cy5.focA-10-30 that pairs to *focA* -30 to -10 region was used to increase the precision of the uRBS toeprints. Orange arrows indicate the uRBS toeprints, and are reported to the uRBS sequence in panel (**a**). **c**. Multiple sequence alignment of *focA* 5'UTR in the Enterobacteriaceae species *Cronobacter sakazakii* CS-931 (*C.sak*), *Enterobacter clocae clocae* ATCC 13047 (*E.clo*), *Citrobacter freundii* CFNIH1 (*C.fre*), *Salmonella enterica* serovar Typhi CT18 (*S.ent*), *E.coli* K12 MG1655 (*E.col*), and *Pseudocitrobacter faecalis* CCM8478 (*P.fae*). The uRBS and the uSD motif are framed, and the *E.coli* TSS is indicated by an arrow. * : Other known *focA* TSSs (positions -27 and -37 relative to Start).
